# Assessing Combinability of Phylogenomic Data using Bayes Factors

**DOI:** 10.1101/250969

**Authors:** Suman Neupane, Karolina Fučíková, Louise A. Lewis, Lynn Kuo, Ming-Hui Chen, Paul O. Lewis

## Abstract

With the rapid reduction in sequencing costs of high-throughput genomic data, it has become commonplace to use hundreds of genes/sites to infer phylogeny of any study system. While sampling large number of genes has given us a tremendous opportunity to uncover previously unknown relationships and improve phylogenetic resolution, it also presents us with new challenges when the phylogenetic signal is confused by differences in the evolutionary histories of sampled genes. Given the addition of accurate marginal likelihood estimation methods into popular Bayesian software programs, it is natural to consider using the Bayes Factor (BF) to compare different partition models in which genes within any given partition subset share both tree topology and edge lengths. We explore using marginal likelihood to assess data subset combinability when data subsets have varying levels of phylogenetic discordance due to deep coalescence events among genes (simulated within a species tree), and compare the results with our recently-described phylogenetic informational dissonance index (D) estimated for each data set. BF effectively detects phylogenetic incongruence, and provides a way to assess the statistical significance of D values. We discuss methods for calibrating BFs, and use calibrated BFs to assess data combinability using an empirical data set comprising 56 plastid genes from green algae order Volvocales.

## INTRODUCTION

Until recently, common practice for inferring multi-gene phylogenies involved concatenation of all available genes with an assumption that the evolutionary histories of all sampled genes were identical. However, phylogenetic trees for different genes (gene trees) can differ from each other, from the tree inferred from the concatenated data, and from the true species tree, due to evolutionary events/processes such as incomplete lineage sorting (ILS), horizontal transfer, and hybridization (Maddison, 1997; Edwards, 2009; Degnan and Rosenberg, 2009; Mallet et al., 2016). Further, even if the sampled genes share the same evolutionary history, estimated trees can differ because of: (1) insufficient phylogenetic information in the sampled genes (stochastic or sampling error), or (2) model misspecification (systematic error) leading to, for example, long edge attraction in some gene trees and not in others (Swofford et al., 1996; Philippe et al., 2005, Philippe et al.,2011).

With the recent surge of large-scale genomic DNA data from high-throughput sequencing methods, the issue of phylogenetic incongruence has become even more important in phylogeny reconstruction. Inferring species trees by addressing these challenges has become an area of active research in phylogenetics. Several species tree methods already available (reviewed in Liu et al., 2015) are effective in correcting incongruences due to deep coalescence (e.g. Song et al., 2012; Xi et al., 2014; Jarvis et al., 2014; Tang et al., 2015). These methods estimate a species tree either from multiple sequence alignments (e.g. *BEAST, Heled and Drummond, 2010; BEST, Liu et al., 2008; SVDquartets, Chifman and Kubatko, 2014, 2015) or summary statistics calculated from estimated gene trees (e.g. STEM, Kubatko et al., 2009; MP-EST, Liu et al., 2010; BUCKy, Ané et al., 2007; ASTRAL, Mirarab et al., 2014b). Methods such as *BEAST and BEST simultaneously estimate gene trees and the species tree by using MCMC to integrate over trees and substitution model parameters; however, co-estimation of species and gene trees under a multispecies coalescent model is computationally intensive and cannot be applied to large scale genomic data. On the other hand, fast and efficient summary statistic methods (e.g. Mirarab et al., 2016) that completely rely on the estimated gene tree/trees (partial data) for the downstream species tree estimation may be prone to systematic bias as they do not incorporate uncertainty in the gene tree estimation process. Still lacking is a comprehensive approach that employs both a rigorous and more efficient algorithm to estimate species trees with high accuracy from hundreds of loci by addressing not just one (e.g. ILS) but all sources of phylogenetic incongruence (Posada, 2016). Until such methods are widely available, there is a need to at least identify phylogenetically congruent sets of loci among sampled genes. Phylogenies from congruent sets of genes may then be used to estimate a species phylogeny (cf. statistical binning, Mirarab et al., 2014a). Furthermore, identifying genes that are significantly incongruent may also be used to identify sequences resulting from processes other than the standard vertical inheritance model assumed in most phylogenetic analyses.

### Phylogenetic dissonance

Lewis et al. (2016) introduced Bayesian methods for measuring the phylogenetic information content of data and for measuring the degree of phylogenetic informational dissonance among data subsets. Phylogenetic dissonance is relevant to the problem of identifying congruent subsets of loci. When data are partitioned into subsets(corresponding to, for example, genes or codon positions), such tools yield insight into which data subsets have the greatest potential for producing well supported estimates of phylogeny. Conflict between different subsets with respect to tree topology can lead to paradoxical results with respect to both information content and estimated phylogeny. For example, a tree topology minimally supported by all subsets (posterior probability less than 0.2) may be given maximal support (posterior probability 1.0) in a concatenated analysis if each subset is highly informative and effectively rules out the trees most supported by other subsets (Lewis et al., 2016). The information measure *D* (phylogenetic dissonance) was introduced by Lewis et al. (2016) to specifically identify such anomalies. Phylogenetic dissonance is defined as

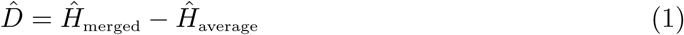

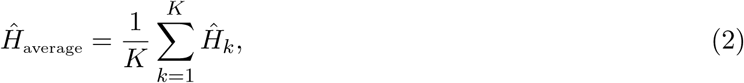

where 
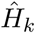
 is the entropy of the marginal tree topology posterior distribution for data subset *k* (of *K* subsets), and 
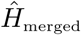
is the entropy of a posterior distribution estimated from a merged tree sample. Posterior tree samples from separate analyses of each data subset are combined to form the merged tree sample. (Note that this merged tree sample differs from a tree sample obtained from a concatenated analysis.) If different data subsets strongly support mutually exclusive tree topologies, then the average entropy of marginal tree topology posterior distributions 
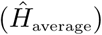
 will be small while the merged entropy 
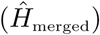
will be relatively large due to the fact that topology frequencies are more evenly distributed in the merged sample compared to samples from individual subsets, which are each dominated by one tree topology. Lewis et al. (2016) defined and estimated phylogenetic dissonance using this entropy-based measure, but how to evaluate the statistical significance of a given level of phylogenetic dissonance remains an open question.

### Tests for Phylogenetic Dissonance

The only direct tests of phylogenetic congruence proposed to date are likelihood ratio tests (LRTs). Huelsenbeck and Bull (1996) proposed a parametric bootstrapping approach in which the null hypothesis constrained all data subsets to have the same tree topology, while the alternative (unconstrained) hypothesis allowed each subset to have a potentially different tree topology. The distribution of the test statistic was generated by simulating data sets under the null hypothesis using maximum likelihood estimates of all model parameters and computing the test statistic under each simulated data set.

Non-parametric boostrapping, in conjunction with LRTs, was used by Leigh et al. (2008) to test the same null hypothesis. Leigh et al. (2008) also proposed clustering of data subsets based on pairwise LRT results to generate compatible sets. Separate likelihood ratio tests were also proposed by Leigh et al. (2008) to test for heterotachy: in this case the null hypothesis constrains edge lengths to be proportionally identical across subsets, while the alternative hypothesis allows each subset to potentially have different edge lengths. The software CONCATERPILLAR (Leigh et al., 2008) may be used to carry out these non-parametric bootstrapping LRTs.

These likelihood ratio tests are well justified and are the best available means to assess congruence when there are no priors involved in the tree estimation process. However, when the phylogeny estimation involves Bayesian methods, then evaluation of congruence should properly account for the effects of the assumed prior distributions. We propose a Bayesian approach to testing phylogenetic congruence (or, equivalently, dissonance) by comparing the marginal likelihoods of competing models. When only two models are compared, the ratio of marginal likelihoods is termed the Bayes Factor (BF). Our approach is comparable to that of Leigh et al. (2008), but instead of comparing maximized log-likelihoods of competing models using LRTs, we use marginal likelihoods and their ratio (BF) for model comparison. Our approach is made possible by the recent improvements in marginal likelihood estimation (stepping-stone, SS: Xie et al., 2010, Fan et al., 2011; path-sampling, PS: Lartillot and Philippe, 2006; partition weighted kernel estimator, PWK: Wang et al., 2017) for phylogenetic model selection. The SS and PS estimators substantially outperformed other approaches (e.g. harmonic mean estimator, HME, and a posterior simulation-based analog of Akaikes information criterion through Markov chain Monte Carlo, AICM) for comparing models of demographic change and relaxed molecular clocks (Baele et al., 2012). Recently, Brown and Thomson (2016) also used BF to analyze the sensitivity in clade resolution to the data types used to infer the topology. The primary aim of our study is to evaluate the effectiveness of BF for assessing significance of the phylogenetic dissonance measure *D* (equation 1). We explore the behavior of BF using simulations designed to create a spectrum of 10-gene data sets ranging from low to high information content and from complete topological concordance to extreme discordance (due to deep coalescence and subsequent incomplete lineage sorting). We also provide an empirical example involving concordance of nuclear and plastid genes in the green algal order Volvocales which demonstrates that likelihood ratio tests carried out using CONCATERPILLAR can differ from conclusions based on marginal likelihoods when analyses are performed in a Bayesian context.

## MATERIALS AND METHODS

### Bayes Factors

In Bayes’ Rule,

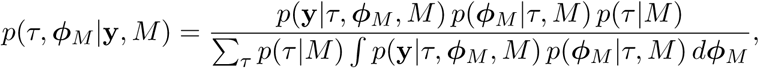

the denominator represents the marginal likelihood *p*(**y**|*M*): the total probability of data **y** given model *M*, averaged over tree topologyand τ a multivariate parameter vector *ϕ*_*M*_ comprising model parameters. The parameters composing *ϕ*_*M*_ may be tree-specific (e.g. edge lengths) or substitution-model-specific (e.g. transition/transversion rate ratio). Data y is a vector comprising observed patterns of states for all taxa for individual characters (sites in the case of sequence data). Considering two models, (*M*_1_, *M*_2_), and their marginal likelihoods, *p*(**y**|*M_1_* and *p*(**y**|*M_2_*, respectively, the **BF** *B*_12_ is the ratio *p*(**y**|*M*_1_)*/*p**(**y**|*M*_2_). The BF on the log-scale is calculated as:

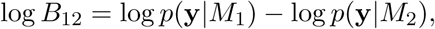

where log *B*_12_ >0 signifies that model *M*_1_ is preferred over *M*_2_. By preferred, we mean that model *M*_1_ fits the data better on average than model *M*_2_ over the parameter- and tree-spacede defined by the prior. Applying this approach to the problem of phylogenetic congruence, consider data from a set of *K* loci y (y_1_, y_2_,…y_*K*_), and two models, CONCATENATED and SEPARATE. The CONCATENATED model represents the marginal likelihood of the concatenated set (y_*C*_) in which all loci are forced to have the same topology and model parameters (*ϕ*_*M*_),

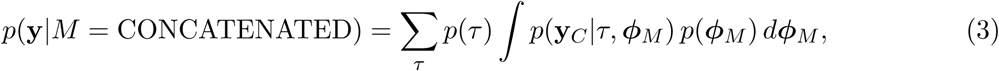

whereas the SEPARATE model represents the marginal likelihood for a model in which individual loci are allowed to have their own topologies and model parameters (*ϕ*_*M*1_, *ϕ* _*M*2_…, *ϕ*_*M K*_),

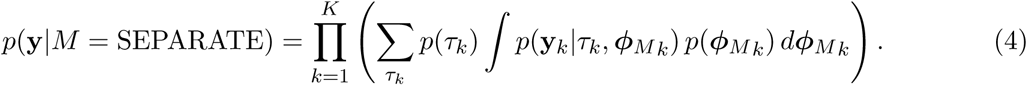

The BF for CONCATENATED against SEPARATE is defined

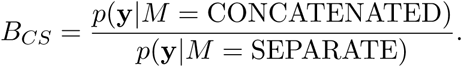

When the tree topology prior is discrete uniform,

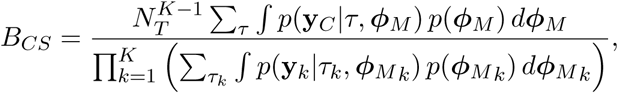

where *N*_*T*_ equals the number of distinct labeled tree. Here, *B*_*CS*_ *>* 1 (or equivalently log *B*_*CS*_*>* 0) indicates that the CONCATENATED model (numerator) is preferred over the SEPARATE model (denominator), whereas *B*_*CS*_ *<* 1 (log *B*_*CS*_ *<* 0) indicates the reverse (i.e. SEPARATE model is the preferred model).

A third, intermediate model HETERO links topology across subsets but allows edge lengths to vary between single-gene data sets:

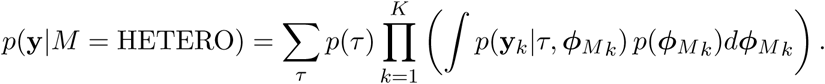

While BF may be de ned between any pair of models, and while we continue to describe our approach as using Bayes Factors, in practice we will only implicitly compute BF, instead estimating the log marginal likelihood of each of the three models and declare the winning model as the one having the largest of the three log marginal likelihood values.

### Data Simulation

Gene trees were simulated within species trees using parameter combinations that yielded differing levels of phylogenetic incongruence. Using a Python script (source code provided in Supplementary Materials), one thousand 6-taxon species trees were generated under a pure-birth (Yule) process in which the tree height *T* (expected number of substitutions along a single path from root to tip) was drawn from a Lognormal(0.05, 0.22) distribution (mean 1.08, 95% of samples between 0.68 and 1.62). Ten gene trees were simulated within each species tree using coalescent parameter *θ*= 4*N*_*eμ*_, where *N*_*e*_ is the effective (diploid) population size and *μ* is the mutation rate per generation. For each species tree, the ratio *θ/T* was drawn from a Lognormal(0.60, 0.77) distribution (which has mean 2.45 with 95% of samples between 0.40 and 8.24) and *θ* was determined by multiplying this ratio by the value of *T* used for a specific species tree. Increasing *θ* relative to *T* results in a higher number of deep coalescences, causing increased discordance among the gene trees.

The gene trees thus generated were subsequently used to simulate DNA sequence alignments of length 2000 sites using seq-gen (Rambaut and Grass, 1997) under the HKY+G model. Individual single-gene datasets and the concatenated dataset were used to compute marginal likelihoods using the Stepping-stone method (Xie et al., 2010) implemented in MrBayes (Ronquist et al., 2012). For the concatenated dataset, two marginal likelihoods were estimated by enforcing: (1) the same topology and edge lengths for all sites (CONCATENATED model), and (2) the same topology but allowing edge lengths to vary among single-gene data subsets to account for non-topological gene tree variation (HETERO model). Analyses of single-gene data sets alone yielded marginal likelihoods that, when multiplied together, yield the marginal likelihood under the SEPARATE model.

In order to assess the robustness of BF for detecting topological and edge length congruence, the BF results were evaluated with respect to the phylogenetic information content (*I*) and phylogenetic dissonance (*D*) values computed using Galax v1.0.0 (Lewis et al., 2016). Estimation of *I* and *D* uses conditional clade probabilities (Larget, 2013) to estimate Shannon entropy (Shannon, 1948), from which 
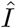
 is calculated simply as a difference between the entropies of the marginal prior and marginal posterior distributions of tree topology (Lindley, 1956). The phylogenetic dissonance is defined as in equation (1), and thus 
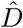
 is computed as the entropy of the merged tree sample minus the average entropy of tree samples from individual genes. We also tested the strength of different variables including 
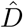
 (and their combinations) in discriminating SINGLE vs. CONCATENATED model by conducting a linear discriminant analysis (LDA). The LDA was carried out in R using the 'lda' function available in the library MASS (Venables and Ripley, 2002) for all the predictor variables (number of conflicting nodes, number of variable sites, number of parsimony informative sites, *θ/T*, species tree height/shortest gene tree height, species tree height/longest gene tree height, average information content, *D*, and number of deep coalescences).

Phylogenetic dissonance is expected to be zero for comparisons of independent MCMC samples from the same posterior distribution, and thus provides a sensitive measure of MCMC convergence with respect to tree topology (Lewis et al., 2016). We replicated each single-gene and concatenated MCMC analysis and computed 
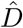
 for these paired samples as a way of ensuring that post burn-in MCMC sample size was sufficient for convergence.

### Lindley’s Paradox

The tendency of Bayes Factors to prefer a sharp null hypothesis (e.g. a point mass prior) over an a priori diffuse (e.g. noninformative) alternative hypothesis when a classical frequentist hypothesis test would reject the null hypothesis is known as Lindley's Paradox (Jeffreys, 1939; Lindley, 1957). The BF is identical to the posterior model odds given equal model prior probabilities. Giving both the sharp null hypothesis and the diffuse alternative hypothesis equal prior weight provides a distinct advantage for the null hypothesis as long as the null hypothesis represents a better explanation of the data compared to most parameter values supported by the alternative hypothesis. The amount of this advantage grows with the a priori diffuseness of the alternative hypothesis.

Consider the BF for CONCATENATED against SEPARATE models. Equation (3) shows that the marginal likelihood of the CONCATENATED model contains a term *p*(*τ*) that equals the prior probability of the tree topology shared among all data subsets. Assuming a discrete uniform prior distribution over tree topologies, *p*(*τ*) is a constant equal to 1/*N*_*T*_. Equation (4) shows that the corresponding term in the marginal likelihood for the SEPARATE model is (1/*N*_*T*_)^*K*^, reflecting the fact that each of the *K* genes potentially has a different tree topology. As either *N*_*T*_ or *K* increases, the CONCATENATED model becomes increasingly sharp compared to the SEPARATE model with respect to prior distributions and thus Lindley’s paradox must be taken into consideration given a sufficiently large number of taxa and/or data subsets. In other words, for large trees or large number of genes, or both, assuming a common tree for all genes may provide a better explanation, even if incorrect in some details, than allowing each gene to have its own tree topology (and independent set of edge lengths). Here, model *fit* is viewed from the Bayesian perspective and is thus more appropriately described as *averaget fit*. is the fact that model fit is averaged over a very large number of incorrect trees, each considered equal by the prior, that drags down the marginal likelihood of the SEPARATE model.

Using BF for testing data combinability must keep the possibility of Lindley's Paradox in mind. Fortunately it is not difficult to determine if Lindley's Paradox applies: if the likelihood ratio test approach chooses the SEPARATE model but BF chooses CONCATENATE, this provides a strong hint that it is the vagueness of the prior in the SEPARATE model that is tipping the balance. While this is less a paradox than a difference in Bayesian vs. Frequentist perspective, a researcher may nevertheless wish to lessen the impact of the tree topology prior on the model choice decision.

While the prior distributions for edge lengths and substitution model parameters are potentially relevant to Lindley's paradox, these parameters are not directly involved in the test and are integrated out of both numerator and denominator in the BF calculation. Bergsten et al. (2013) identified similar issues related to diffuse tree topology priors in BF used for testing monophyly. In that case, constraints placed on tree topologies to enforce monophyly affect the size of tree space, which creates an imbalance in tree topology priors analogous to that encountered when testing for data combinability.

### BF Calibration

It is standard practice to use the value *BF* = 1 as the critical value determining whether the null model (e.g. CONCATENATED) or the alternative model (e.g. SEPARATE) wins. This makes sense when the prior predictive error probabilities of BF under both models are equal; however, in cases where models differ substantially in their effective dimensions, the distributions of the BF for the two models being compared may not be symmetrical. For example, it is possible that the probability of choosing the CONCATENATED model when the SEPARATE model is true may not equal the probability of choosing the SEPARATE model when the CONCATENATED model is true:

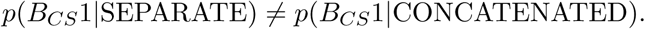

Under such circumstances, a different threshold value (other than 1) can be selected such that the probability of choosing the incorrect model under both hypotheses is equal. Garcia-Donato and Chen (2005) suggested a method for calibrating the BF that makes the prior predictive error probabilities symmetrical. To apply the method of Garcia-Donato and Chen (2005), we simulated 1000 replicate 6-taxon, 10-gene data sets (2000 sites/gene) from the joint prior distribution of each model (CONCATENATED and SEPARATE). For the CONCATENATED model, data for all 10 genes were simulated from a single topology sampled from the discrete uniform topology prior. For the SEPARATE model, data for each of the 10 genes was simulated from topologies separately sampled from the discrete uniform topology prior. All other model parameters were simulated from their respective prior probability distributions.

For each simulated data set, *B*_*CS*_ was computed, yielding a sample of 1000 values from the prior predictive BF distributions for both the CONCATENATED and SEPARATE models. The 2000 sampled BF values were combined into a single vector and sorted, and the critical value *c* was chosen as the midpoint between the 1000th and 1001th values in the sorted vector. This procedure identifies a BF cutoff value *c* that satisfies

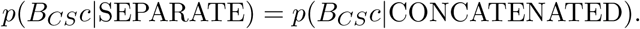

The simulations needed for BF calibration were carried out using PAUP* 4a158 (Swofford, 2003).

### Example from the Green Algal Order Volvocales

We tested phylogenetic congruence among 56 protein-coding plastid genes used in Fučíkováet al. (2016), focusing on one of the most topologically consistent parts of the tree, the green algal order Volvocales. The Volvocales dataset consisted of a subset of the Sphaeropleales, Vovocales, and OCC (Oedogoniales-Chaetopeltidales-Chaetophorales) clades studied in Fučíková et al. (2016). We included four of the five Volvocales members from the study: *Chlamydomonas reinhardtii*, *Gonium pectoral*, *Pleodorina starrii*, and *Volvox carteri*. The length of post-trimmed plastid genes ranged from 93 sites (*psb*T) to 2259 sites (*psa*A). We conducted BF tests for all possible pairs from the 56 genes (by estimating marginal likelihoods under the CONCATENATED and SEPARATE models) used in the study with the aim to detect possible outlier genes that may be present among the sampled genes for the concatenated phylogeny. The critical value *c* for this analysis was computed using the same approach as simulated data. The prior predictive distributions of BF under CONCATENATED and SEPARATE models were obtained from 1000 replicates (4-taxon, 2 genes/replicate, and 2000 sites/gene) simulated under each model using PAUP 4a158 (Swofford, 2003). For the CONCATENATED model, DNA sequence data for both genes were simulated from a single topology (randomly drawn from the discrete uniform topology prior) with edge lengths and other model parameters drawn from the GTR+G model prior distribution, whereas for the SEPARATE model, sequence data for each of the 2 genes were simulated from individually drawn discrete-uniform-distributed topologies with all other model parameters drawn from the GTR+G model prior distribution. In order to compare our results with the likelihood-based approach, we also tested congruence among these 56 genes using CONCATERPILLAR (Leigh et al., 2008) using its topological congruence test (-t) option.

Parameters of the models and the priors used in the study of simulated (model: HKY+G) and Volvocales (model: GTR+G) data were:

Tree topology T ∼ Discrete Uniform(1,*T*)

Tree length *L* ∼ Exponential(0.1)

Edge length proportions e ∼ Dirichlet(1,…,1)

Nucleotide frequencies π ∼ Dirichlet (1, 1, 1, 1)

transition/transversion rate ratio *k* ∼ Beta(1,1)

Exchangeabilities **r** ∼ Dirichlet (1, 1, 1, 1, 1, 1)

Discrete Gamma shape α ∼Exponential(1),

where *T* equals the total number of distinct, labeled, binary, unrooted tree topologies.

## RESULTS

### Bayes Factor Calibration For Simulation Study

BF calibration for the 6-taxon, 10-gene simulation study resulted in a critical value *c* =−3.2 (log scale), which equals the midpoint in the interval extending from the 1000^th^ element (−3.55) to the 1001^st^ element (-2.85) of the combined, sorted vector of prior predictive log *B*_*CS*_ values from CONCATENATED and SEPARATE models. Using the standard log-BF cutoff (0.0) would thus result in the SEPARATE model winning more often when CONCATENATED is the true model than the CONCATENATED model wins when the SEPARATE model is true.

### Phylogenetic Dissonance Correlated with Number of Deep Coalescences

As expected, estimated phylogenetic dissonance 
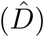
 was correlated with number of deep coalescences in 1000 simulated gene sets (10 genes/set) representing various degrees of topological and edge length congruence (Fig. 2). The number of deep coalescences varied from the minimum possible number (0) to the maximum possible number (50). (The maximum number of deep coalescences is 5 per gene because there are 5 internal nodes in a rooted tree of 6 taxa.)

**Figure 1:**
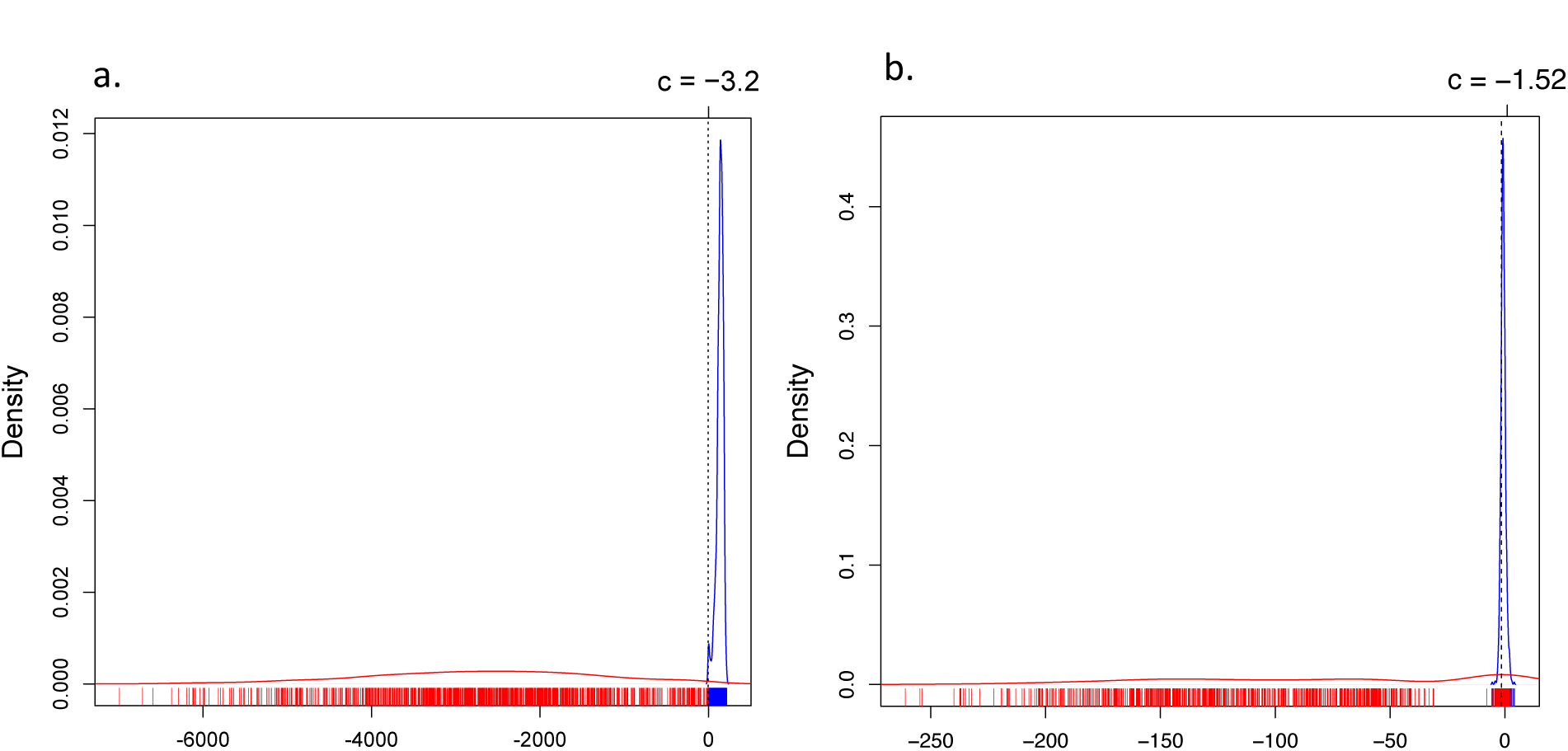
Density of *B_CS_* under CONCATENATED (blue line) and SEPARATE (red line) models for the (a) 6 taxa, 10-gene data set and (b) 4 taxa 2-genes data set. The critical values (*c* = −3.2, *c* = −1.52) are indicated by dashed lines, estimated using 1000 prior predictive replicates from each model (rug).

**Figure 2:**
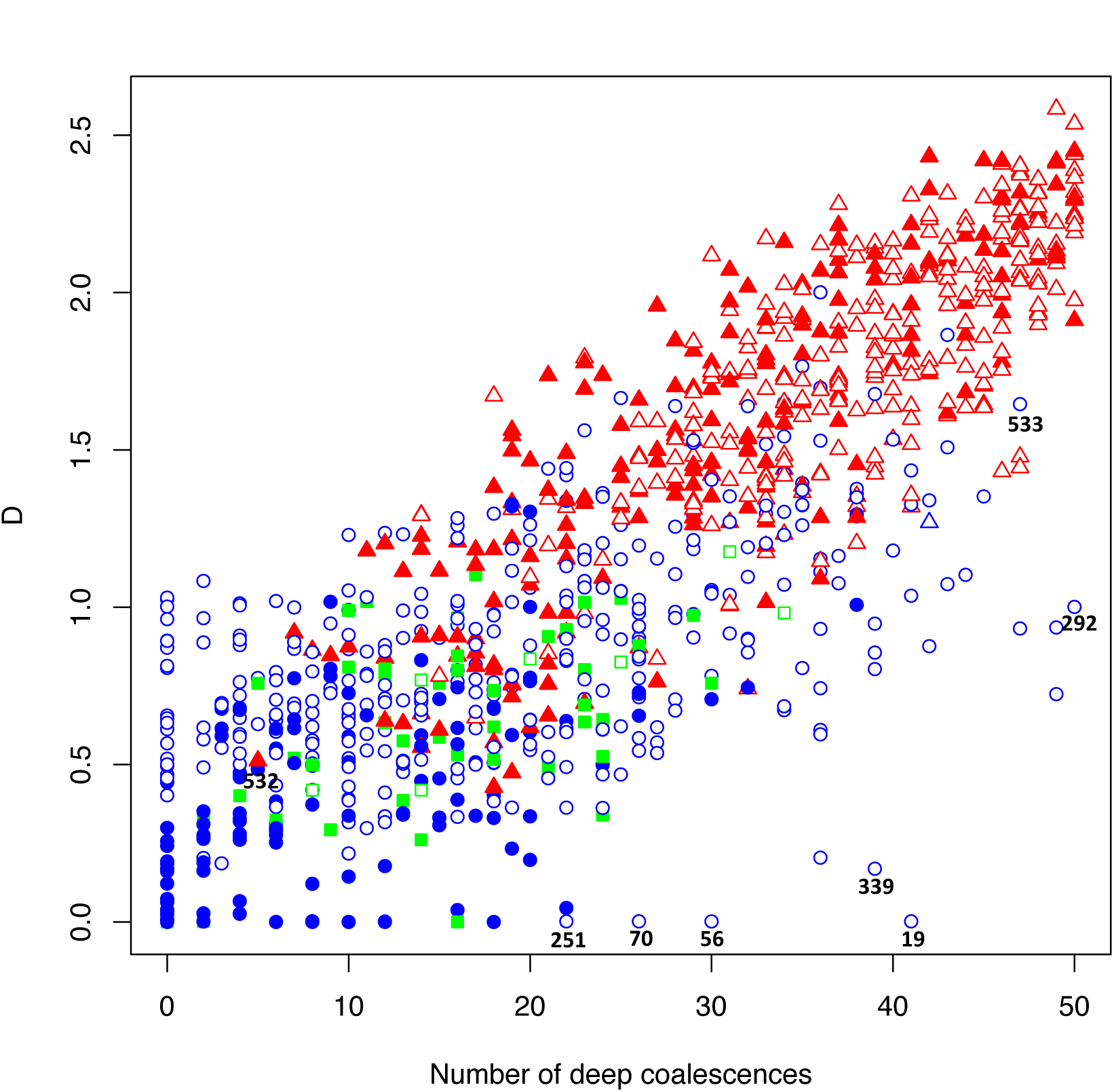
Plot of 1000 replicates simulated under conditions that yielded varying levels of deep coalescence (x-axis) and phylogenetic dissonance 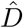 values (y-axis). Blue circles indicate Bayes Factor support for the CONCATENATED model over both HETERO and SEPARATE with log-scale critical value *c* = 0. Blue triangles indicate Bayes Factor support for the CONCATENATED model over both HETERO and SEPARATE with −3:2 *< c <* 0. Green square indicates support for the HETERO model over both CONCATENATED and SEPARATE. Red triangle indicates support for SEPARATE model over both CONCATENATED and HETERO with *c* = −3.2. Filled symbols represent ≥90% average information content across genes, with closed symbols indicating <90%. Numbers indicate particular replicates mentioned in the text.

In our simulations, under both criteria (*c* = 0, *c* = −3.2), the SEPARATE model won in a majority of replicates when 
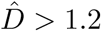
or when the number of deep coalescences exceeded 1.8 per gene. Under the new critical value (*c* = −3.2), 1 simulation replicate switched its support to the CONCATENATED model from the earlier SEPARATE or HETEROTACHY model. When SEPARATE failed to have the largest log marginal likelihood, CONCATENATED usually won, with HETERO only achieving the largest log marginal likelihood if 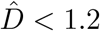
and the number of deep coalescences was less than 3.2 per gene.

In cases of multiple deep coalescences (*>*3.2/gene) or high dissonance 
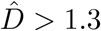
, the CONCATENATED model won only when average information content was low, while HETERO never won under these circumstances. In a few cases, SEPARATE was the winning model even when the number of deep coalescences was less than 1 per gene on average. Conversely, CONCATENATED was occasionally the winner despite high levels of deep coalescence (*>* 3.5 per gene).

### Volvocales Example

The results of the pair-wise tests of congruence are illustrated in Fig. 3a. The critical value *c* computed for the four-taxon case based on the prior predictive distributions of BF under CONCATENATED and SEPARATE models was −1.52 (Fig. 1b). Under both criteria (*c* = 0, *c* = −1.52), marginal likelihoods indicated congruence for all gene pairs with the exception of *pet*D and *rpl*36, each of which was incongruent with every other gene (but were congruent with each other). Both *pet*D and *rpl*36 favor *Gonium* + *Pleodorina* (Fig. 3b) while all other genes favor *Volvox* + *Pleodorina* (Fig. 3c). The CONCATERPILLAR analysis, however, indicated that all 56 genes were topologically congruent. The two genes found to be incongruent using BF analysis (*pet*D and *rpl*36) were not contiguous in the chloroplast genomes of four taxa, suggesting that they were not the result of a single horizontal transfer event. In the case of *pet*D, there is a single variable amino-acid site (amino-acid position 106) that determines the *Gonium* + *Pleodorina* relationship. Excluding site 106 removes support for this relationship. Despite the incongruence of *rpl*36 to the other genes, this particular gene is short (total nucleotide length =114) and it contains relatively less information relevant to estimating topology. We also used PhyloBayes (Lartillot et al., 2009) to estimate phylogeny for the *pet*D data (including all the sequences from Chlorophyceae available on Dryad: http://dx.doi.org/10.5061/dryad.q8n0v) under the CAT model (Lartillot and Philippe, 2004). The CAT model accommodates sites with distinct state frequency profiles, unlike standard models that assume state frequencies are homogeneous across sites. The CAT model can potentially avoid long-branch attraction due to the model assuming a wider range of available amino acids at particular sites than are actually available. However, even under the CAT model, *Pleodorina* resolved sister to *Gonium*.

**Figure 3.**
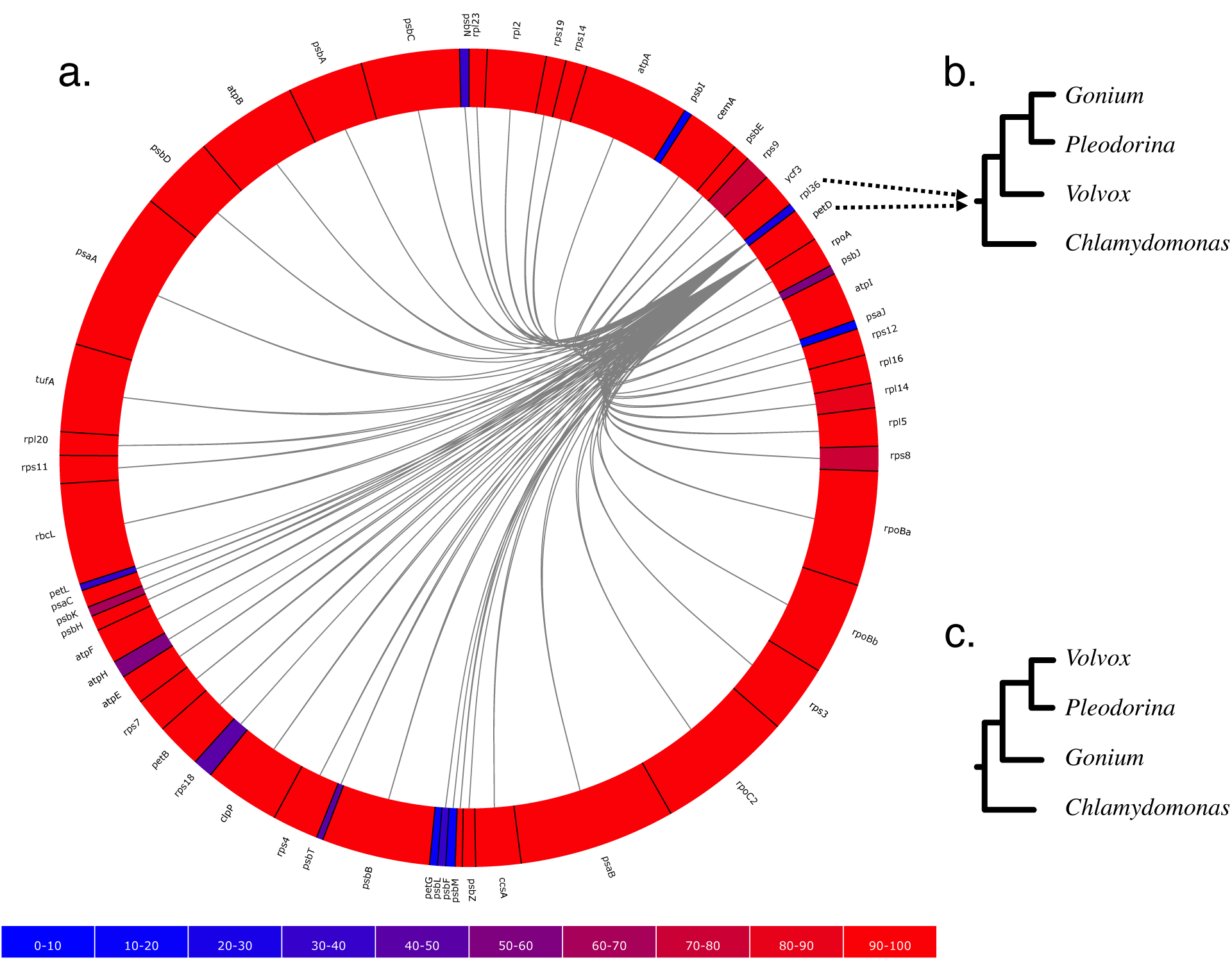
:Pair-wise BF test for phylogenetic congruence for all possible pairs among 56 protein coding plastid genes (Fučíková et al., 2016) where the colors represent the information content present in the gene and the lines between the genes indicate phylogenetic incongruence (i.e. support for SEPARATE over CONCATENATED) suggested by the BF test (3a). In the 56 gene sets, *pet*D and *rpl*36 show support for *Gonium* + *Pleodorina* relationship (3b), whereas the other 54 genes support *Volvox* + *Pleodorina* relationship (3c).

## DISCUSSION

The presence of deep coalescence does not guarantee that different genes will have different tree topologies, but the fact that lineages are joined randomly when there is deep coalescence means that greater incongruence is the expected result of increasing the frequency of deep coalescence. In general, more deep coalescences yielded higher 
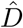
 and a greater chance of the SEPARATE model winning. In fact, *D* was the most important variable in discriminating SINGLE vs. CONCATENATED model in the discriminant function analysis involving a number of other variables tested (number of conflicting nodes, number of variable sites, number of parsimony informative sites, *θ/T*, species tree height/shortest gene tree height, species tree height/longest gene tree height, average information content, *D*, and number of deep coalescences). The *D*, number of deep coalescences, and *θ/T* could separate SINGLE vs. CONCATENATED models with 91% accuracy where the *D* alone could separate the two with 82% accuracy. Because a single simulation study can only suggest appropriate cutoff values for 
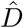
 for the limited range of parameter combinations explored, we argue that a BF approach provides a sensible general approach for determining when values of 
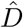
 are too high to be compatible with phylogenetic congruence.

It is interesting and informative to examine some outliers in the simulation results presented in Figure 2. For example, consider replicate 532, for which the SEPARATE model won despite a high average information content (94% of maximum information), relatively low *D*, and a single topological conflict among 10 genes. Removing the gene that conflicted (gene2) from the concatenated set, followed by re-estimation of marginal likelihoods, resulted in a win for the CONCATENATED model, suggesting that a single incongruent subset out of 10 total can be enough to place the SEPARATE model on top.

Low phylogenetic signal can result in a preference for the CONCATENATED model despite a high number of deep coalescences (e.g. Fig. 2, replicates 19, 56, 70, 97, 251, 292, 339, 533). In some extreme cases, when phylogenetic information content is very low (approaching zero information), 
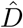
 can also be low (Fig. 2, replicates 19, 56, 70, 251). In such cases, posterior samples of individual gene subsets visit every possible tree topology (of the 105 possible unrooted binary tree topologies for 6 taxa) in roughly equal proportions. Phylogenetic dissonance is zero if all subset posterior distributions are equal, and this is true whether these posterior distributions are concentrated or flat, so low 
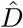
 in the face of low information content for all gene subsets is not surprising. It is also unsurprising that the marginal likelihood would favor the CONCATENATED model in such cases because one tree topology is about as good as any other tree topology in explaining the data, and the marginal likelihood implicitly punishes models for including parameters that do not provide access to regions of parameter space providing appreciably higher likelihood.

### The Case of Mistaken Heterotachy

An interesting phenomenon was observed as a result of using phylogenetic dissonance to assess MCMC convergence with respect to tree topology. Most replicate MCMC analyses exhibited 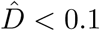
, indicating that the posterior samples from replicate analyses were essentially identical (as they should be if both Markov chains mixed well and were sampled only after converging to the stationary distribution); however, many analyses exploring the HETERO model produced unexpectedly high replicate phylogenetic dissonance values. The reason for this turns out to be the completely understandable result of a model making the best of a bad situation, and offers a warning for those who might be tempted to use a HETERO model win as an evidence for heterotachy.

Consider a case of two data subsets (genes) in which the true tree topology differs (Fig. 4). The HETERO model assumes that the same tree topology applies to both genes (which is *not* true in this case), but allows each gene to have its own set of edge lengths. The HETERO model can choose to focus on the true tree topology for gene 1 and attempt, using edge lengths, to explain the data for gene 2 as best it can. Alternatively, it can focus on the true tree topology for gene 2 and attempt, using edge lengths, to explain the data for gene 1 to the extent possible. How does a model fit data when assuming an incorrect tree topology? The answer is that it increases the lengths of edges for some taxa that are sister taxa in truth but not in the assumed tree, leaving other closely related taxa connected by relatively short paths. Thus, the fact that some taxa are more similar than the tree topology suggests can be explained by the model using evolutionary convergence (the long edged taxa), while similarities between other taxa that seem far apart on the assumed tree topology are explained by a lowered rate of substitution. In replicate analyses, it is possible for one run to choose the tree topology for gene 1 and the other replicate to choose the tree topology for gene 2, yielding posterior distributions that are concentrated on conflicting tree topologies, which in turn produces high estimated phylogenetic dissonance. The lesson to be learned from this study is that a win by the HETERO model may not mean the presence of heterotachy in data, but may simply reflect a model doing its best to explain data generated on a different tree topology. This crafty use of spurious edge lengths by models to explain away topological discordance among genes was explored in detail by Mendes and Hahn (2016). In their study of simulated and empirical data, Mendes and Hahn (2016) found that the topological discordance between gene trees due to ILS can cause multiple apparent substitutions on the focal tree (e.g. species tree) on one or more of its branches that uniquely define a split on the discordant gene tree that is absent in the species/focal tree. It is interesting that measuring phylogenetic dissonance among replicate analyses under the CONCATENATED model alone can potentially be used to detect incongruence in gene tree topologies.

**Figure 4:**
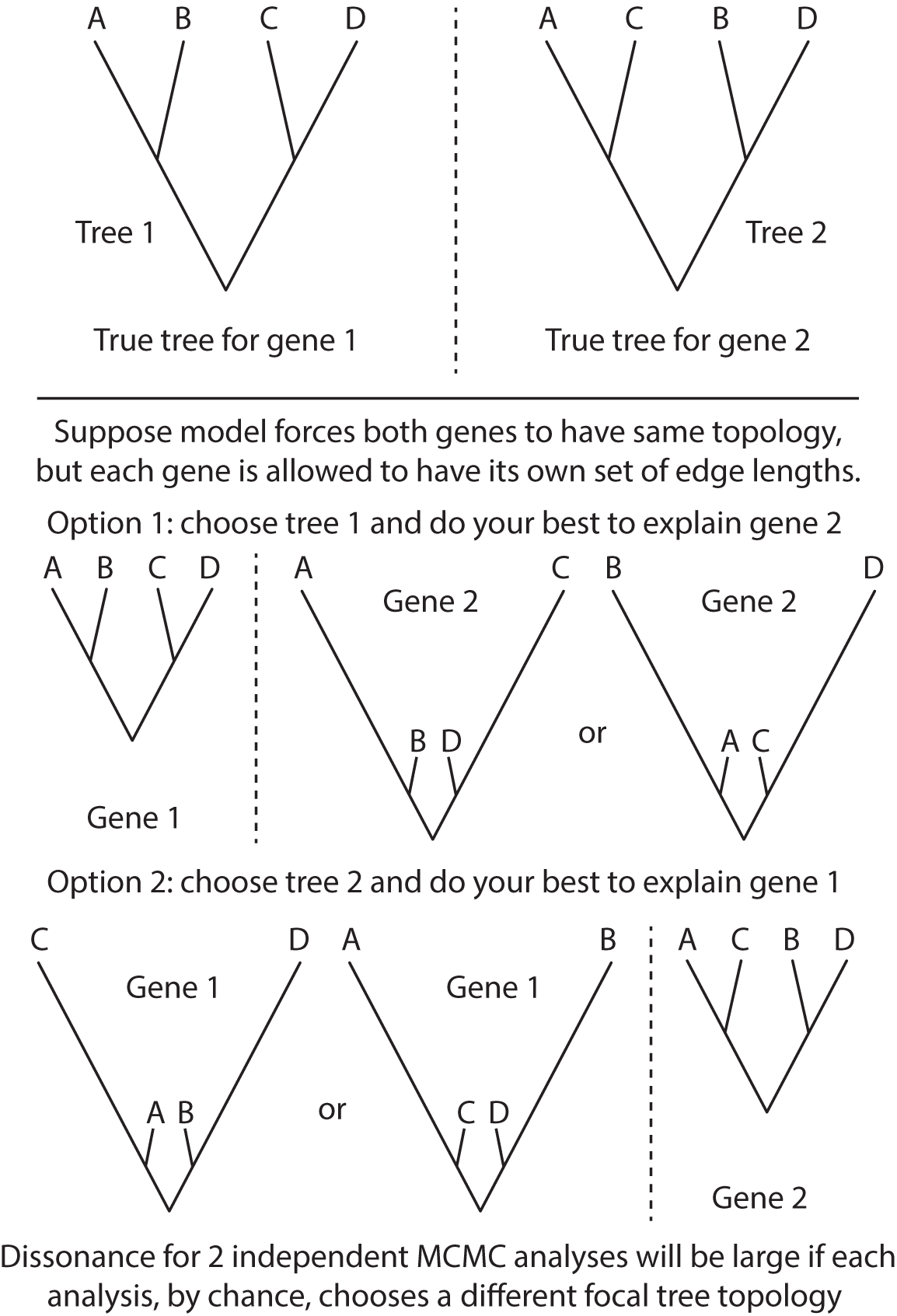
Explanation of paradoxical high dissonance for samples from independent replicate MCMC analyses exploring the same posterior distribution under the HETERO model.

The presence of true heterotachy is suggested by low phylogenetic dissonance combined with HETERO model being the winning model. None of our simulations imparted true heterotachy; however, some results (e.g. replicate 942) did combine 
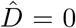
 with a winning HETERO model. The explanation is that the HETERO model is actually detecting *heterochrony* (a new term) rather than *heterotachy*. Heterochrony may be defined as differences in the same edge length (measured in expected number of substitutions per site) across genes due to the fact that coalescence depth varies among genes (even if the topology is identical). The HETERO model is, in this case, detecting differences in coalescence times instead of differences in rate of substitution.

### Empirical Volvocales Example

Our empirical example involved a reanalysis of a subset of four taxa from a more inclusive study of green algal phylogeny. In that former study, Fučíková et al., 2016 found strong support for a single tree topology relating these four taxa using a concatenated dataset, but reported very low internode certainty (IC: Salichos et al., 2014) values for all but one edge in the estimated tree. This suggests some conflict exists among genes, and thus it is not surprising that our BF analyses identified two genes (*rpl*36 and *pet*D) that preferred a different tree topology than the majority (54/56) of genes. What is perhaps surprising is that likelihood ratio tests conducted using CONCATERPILLAR found no conflict, concluding that all 56 genes should be concatenated. The fact that our BF approach and CONCATERPILLAR's LRT approach provide conflicting advice highlights a major difference between the Bayesian and frequentist statistical approaches to phylogenetics. For the *pet*D gene, we found that a single amino acid site (site 106) determines the preference of this gene for *Gonium* + *Pleodorina*. Bayesian analyses do not take into consideration (either implicitly or explicitly) any data other than what was observed, and thus will take the evidence from site 106 at face value. Assuming a site appears (to the model) to be a reliable reporter (i.e. substitution is rare and the site is not contradicted by any other site), then even one site may have a strong impact on a Bayesian phylogenetic analysis. Frequentist approaches involving bootstrapping, however, take additional sources of uncertainty into consideration. Bootstrapping evaluates many data sets, each different than the observed data set, and thus takes uncertainty in the observed data into account. This is one explanation for why bootstrap support values for clades tend to be smaller than posterior probabilities: the Bayesian analysis assumes that there is no uncertainty in the observed data and never considers the possibility that the observed data could be atypical in some way. If support for a clade depends critically on a single site, then the bootstrap support for that clade depends on the probability that the site will be included at least once in a particular bootstrap replicate. The probability *q* that a particular site (out of *n* total sites) will be omitted from any given bootstrap data set is

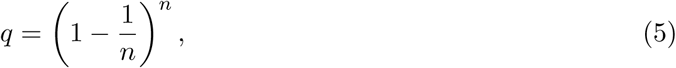

which (by definition) approaches *e*^−1^ as *n* → ∞. Thus, the probability that a single critical site will be included at least once in any given bootstrap data set is *p* = 1 − *q*, which is approximately 63% for a reasonably large number of sites. We should therefore not expect strong bootstrap support for a clade if that clade is supported only by a single site, even if that site appears to be reliable indicator of history. Such a site may, however, have a strong impact on a Bayesian analysis because data sets excluding that site are never considered. For this reason, frequentist tests of data combinability that use bootstrapping to evaluate the significance of likelihood ratios are not appropriate when Bayesian approaches are used for estimating phylogeny.

### SUMMARY

Marginal likelihoods provide a straightforward way of assessing the statistical significance of phylogenetic dissonance (Lewis et al., 2016). We simulated data sets with varying levels of deep coalescence and found, as expected, that larger numbers of deep coalescence events led to higher estimated phylogenetic dissonance and also to preference for the SEPARATE model over the CONCATENATED and HETERO models based on estimated marginal likelihoods. Exceptions mainly involved data sets with low information content due to small tree lengths, which can show low dissonance and preference for the CONCATENATED model despite a relatively large number of deep coalescence events. We calibrated BF comparisons between CONCATENATED and SEPARATE using the method of Garcia-Donato and Chen (2005) to determine the critical value that balances the prior predictive error probabilities of competing models, finding that the standard cutoff (1.0, or 0.0 on the log scale) is not always ideal but in practice changed very few of our model choice determinations. Our results also show that conflict among gene tree topologies may masquerade as heterotachy in combined analyses, as shown previously by Mendes and Hahn (2016).

## ACKNOWLEDGEMENTS

This study benefited from computing resources made available through the computing cluster provided by the Computational Biology Core of the UConn Institute for Systems Genomics. We thank system adminstrator Jeffrey Lary and director Dr. Jill Wegrzyn for their assistance with these computing resources.

